# The Response of Bacterial Flagellar Motor to Stepwise Increase in NaCl Concentration

**DOI:** 10.1101/2020.08.19.257477

**Authors:** V. Soman, S. Kumari, S. Nath, R. Elangovan

## Abstract

Many species of bacteria use flagella to navigate in its environment. The flagellum is a 7-10 μm long helical filament with a rotary motor at its base embedded in the cell membrane and almost a dozen stator complexes. Proton motive force across the cell membrane powers the flagellar motors of *E*.*coli* and *Salmonella*. The motor stochastically switches between clockwise and counter-clockwise direction. A chemotaxis system causes the motor to change its direction, but the process is more complex as the switch is sensitive to load and proton motive force as well. NaCl is significant with regard to the flagellar motor as it affects the stator dynamics, proton motive force, and osmotaxis at higher concentration. Chemotaxis helps the bacteria for its growth and survival. *E*.*coli*’s natural habitat has high osmolarity and the organism uses use various mechanisms for osmoregulation. However, the role of flagellar motor to adapt to the changes in osmolarity, or osmotaxis, is not well studied. In this work, we dissipated the membrane potential of bacteria in pH 7 using step-wise increase in concentration of NaCl in motility buffer and studied the output of E.coli’s flagellar motor using tethered bead assay and swimming Salmonella enteritidis cells. We observed decrease in motor speed and switching rates with stepwise increase in NaCl concentration in the motility buffer. The mean speed of the motors decreased with NaCl concentration. The population of swimming cells tumbled more with increase in concentration of NaCl. At the single motor level, the motors biased to CCW rotation with decrease in membrane potential. In this study, we present our observations of the flagellar motor in high NaCl concentration, and explore how NaCl can be used to study various aspects of the bacterial flagellar motor.

**Statement of significance:** Sodium ion has been significant in the both the cellular energetics and the function of bacterial flagellar motor. Growing evidence show that the effect of sodium ions was not what hitherto thought it would be. It is involved in the sodium energetics, dissipate membrane potential, affect the flagellar stator dynamics of bacteria. Being an osmolyte, it influences the osmotaxis of bacteria. In this work, we studied the effect of NaCl on the response of the single bacterial flagellar motor of *E*.*coli* and swimming cells of *Salmonella enteritidis*. We observed that the effect of NaCl on the output of the flagellar motor was significant and it may affect the cells in various ways.

## Introduction

Bacterial flagellar motor aids in locomotion, navigation, pathogenesis(1, 2), and colonisation on surfaces. The flagellum consists of a rotary unit and stator complex inside the cell, a helical filament that extends from the cell membrane, and a hook that connects the filament with the rotor. The rotor is a supramolecular assembly of proteins, and the stators are ion channels. In *E*.*coli* each stator complex consists of MotA and MotB proteins arranged in a 4MotA-2MotB stoichiometry. The stator complexes are dynamic and their incorporation into the motor depends on the load experienced by the motor(3, 4). The electrochemical energy stored in ion motive force across the cell membrane powers the motor(5–11). The ion transit through the motor converts this energy to the mechanical motion of the flagella; however, the exact mechanism of this energy coupling is unknown. Flagellar motors of *E*.*coli* and *Salmonella* use H^+^ ions (H^+^ type motor) and *Vibrio* species use Na^+^ ions (Na^+^ type motor). The proton motive force (henceforth referred to as pmf) has two components: the electrical potential across the membrane (ΔΨ), and the chemical gradient of protons across the membrane (ΔpH) (12). The proton motive force is calculated as: pmf = ΔΨ+60ΔpH, where ΔΨ is the electrical potential difference between inside and outside of the membrane and ΔpH is the difference in pH between inside and outside of the membrane: 2.3RT/F is ∼60mV at 30°C, where R is the gas constant, T is the absolute temperature, and F is the Faraday constant.

Both the components of pmf contribute equally to the swimming velocity of cells(13), and the speed of the motor changes linearly with the pmf (14). To study the mechanism of chemical coupling with the mechanical motion of the flagella, it is imperative to manipulate the pmf. It has been manipulated by the application of electric pulses, addition of weak acids, and ionophores, and changing intensity of light (11, 13, 15–20). Besides powering the motor, the proton motive force influence stator dynamics(20–23) and the motor switch(24).

The reversibility of motor entails the bacteria to change its swimming direction to evade repellents and to seek attractants. This behaviour is called chemotaxis(25, 26), which includes membrane receptors and a signalling cascade downstream to the receptors (for a review, see(27)). The significance of studying chemotaxis includes understanding the mechanism of flagellar motor and how bacteria survives in their environment. Bacterial membrane has receptors for most chemicals, but some chemoeffectors such as pH and osmolarity has no receptors on the membrane although they elicit a chemotactic behaviour(28).

Osmotaxis is the movement of motile bacteria towards an optimal water content(29–32). Osmotic upshock (increase in osmolarity) and downshock (decrease in osmolarity) causes cell plasmolysis and cell lysis respectively(33, 34). Osmoregulation maintains an optimal turgor pressure which is essential for maintaining growth and cell shape(35). Bacteria adopt several strategies for osmoregulation such as production of osmoprotectants in cytoplasm(36) and periplasm(37), regulating K^+^ transport(38), regulating synthesis of porin proteins(39). Berg has showed that the osmotaxis is due to the perturbation of chemoreceptor due to the changes in cell’s turgor pressure(40). The behaviour of osmolytes varies with their permeability. Osmolyte such as sucrose is slowly permeable(41) and NaCl is highly permeable(42). Rosko studied the effect of sucrose at single motor and population level, and they observed increase in motor speed with external osmolarity and the motor is biased to clockwise direction in high osmolarity solutions (repellent-like response)(31). At the single cell level, less is known about the behaviour of flagellar motor in the presence of a highly permeable osmolyte like NaCl. The membrane potential of bacteria is dissipated in the presence of NaCl(43), and the presence of Na+ ions may affect the stator dynamics(20, 44). Hence, the presence of NaCl in the external environment affect the bacteria in several ways, which makes it an interesting study.

In this paper, we dissipated the membrane potential of bacteria with NaCl in pH 7, and performed temporal assay at single motor level to population level by changing the concentration of NaCl in motility buffer to study the single motor of *E*.*coli* and swimming behaviour of *Salmonella enteritidis*. We increased the concentration of NaCl stepwise until the motors stopped. We found that the increase in concentration of NaCl decreased the motor’s speed (and swimming speed of cells) and changed the rotational bias of single motors.

## Materials and Methods

### Bacterial strains

For swimming assay, *Salmonella enteritidis* MTCC 3219, which is wild type for motility, was used. It is a rod shaped, peritrichously flagellated, facultative aerobic, gram-negative bacterium. For membrane potential measurement and tethered bead assay, *E. coli KAF84* (a generous gift from Howard Berg’s lab) was used. It is wildtype for chemotaxis and has a plasmid with ampicillin resistance and a *fliC*^*sticky*^ gene, which produces flagellar filaments that stick readily to glass surfaces and hydrophobic beads.

### Culturing and growth of strains

Both the strains were grown on Tryptone broth (1% Bacto tryptone, 0.5% NaCl). A single colony *Salmonella enteritidis* MTCC 3219 was sub-cultured in 10 ml tryptone broth media at 37°C and shaken at 200 rpm until the cells reached the mid-exponential phase (0.4-0.6 OD). The cells were washed by centrifugation at 3200 rpm for 4 mins with pH 7 motility buffer, which was 10 mM KPi, 0.01 mM EDTA, and 0.002% Tween 20, and resuspended in fresh buffer. EDTA was added to chelate divalent ions that prevent motility and Tween 20 was to prevent clumping of cells. The motility buffer without NaCl is henceforth referred to as MB^-^, motility buffer with 70 mM NaCl as MB, motility buffer with NaCl concentration more than 70 mM is MB^+^. For *E*.*coli* KAF84 strain, the tryptone broth was supplemented with 100 μg/ml ampicillin. The culture was grown at 33 °C while being shaken at 200 rpm. The cells were harvested when the OD_600_ reached between 0.6 - 0.8. The culture was washed twice with MB^-^ and resuspended in fresh motility buffer.

### Bright field motility assay

#### Sample preparation and image acquisition

A flow chamber was made by layering a square coverslip (22 mm x 22 mm) on a rectangular glass slide (60 mm x 24 mm) with a double sticky tape. The volume of the flow chamber was ∼30 μl. Before image acquisition, the cells were suspended in MB-was washed twice and resuspended in fresh motility buffer with appropriate NaCl concentration. Just after resuspension, about ∼30 μl of cell suspension was added to the flow chamber, and the slide was mounted on a custom-built temperature controller maintained at 30°C on the microscope. Olympus IX 71 inverted microscope equipped with a 1.4 NA 60X oil immersion objective, sCMOS camera (Andor Zyla), and halogen lamp for bright field illumination were used. A time series of bright field images were captured with 30 ms exposure time for 100 frames (∼20 frames/s). The image area was 1392×1040 pixels and the size of the pixel was 0.122 μm. The movies were captured using ImageJ Micro Manager camera software.

#### Image processing and data analysis

The grayscale brightfield images were manually converted into binary images by thresholding in ImageJ software. The files in each movie were named as per the requirements of the analysis program and saved separately. TumbleScore analysis program was used to quantify the motility of *Salmonella enteritidis* in various NaCl concentration. It is a Matlab based program to process bacterial motility movies to obtain swimming behaviour metrics(45). It provided metrics such as linear speed, RCD (rate of change of direction), and Arc-to-chord ratio (ACR) which provides information on tumbling of cells. This program was automated in selecting the tracks, which avoids bias, and it provided these metrics with a low-frame rate motility video. For more details of the program, see reference(45)).

Movement of z-plane was not tracked by the program. Cells with z-component had disjointed trajectories and were thus eliminated. So, it computed only the movement of cells in the x-y plane. The program reported trajectories of cells in each movie, and the velocity was calculated from the trajectories and averaged.

### Measurement of membrane potential (ΔΨ)

The carbocyanine dye DiOC_2_(3) (3,3′-diethyloxacarbocyanine iodide) was used to measure the membrane potential of *E*.*coli KAF 84* cells in pH 7. The dye was a part of BacLight bacterial membrane potential kit from ThermoFisher Scientific®. It is a lipophilic cationic dye which enters the bacterial cells as monomer (emits green fluorescence) and self-aggregates in the cytoplasm. The monomer emits green fluorescence (520nm) and the aggregated dye emits red fluorescence (620 nm). The aggregation depends on the size of the membrane potential. The red fluorescence depends on both the size of the cell and the membrane potential, and the green fluorescence depends only on the size of the cell. Hence, the membrane potential of bacterial cells can be quantified by measuring the Red/Green (R/G) fluorescence ratio as the ratio eliminates the size difference among cells. Victor Nivo Multiplate reader was used to measure fluorescence. Cells suspended in pH 7 motility buffer were washed twice and resuspended in 50 mM pH 7 HEPES buffer, 1 mM EDTA for 10 minutes to permeabilise the cell membrane for the dye. Next, the suspension was washed twice and resuspended in 50mM pH 7 HEPES buffer. Standard curve of Red/Green fluorescence ratio with the membrane potential of the cell membranes was made as follows: The reaction volume was 300 μl. The external concentration of potassium required to induce an external potential was calculated based on the Nernst equation and table below:

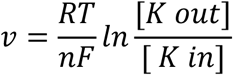

|  |  |  |  |  |  |
| --- | --- | --- | --- | --- | --- |
| 0mV | -25mV | -50mV | -75mV | -100mV | -125mV |
| 300mM | 113.4mM | 46mM | 18.1mM | 7.1mM | 2.3mM |

(assuming cellular potassium concentration as 300mM)

150 μl of cells suspended in HEPES buffer were added to the 6 wells corresponding to the concentration of potassium as shown above. From 1M KCl stock solution (in HEPES buffer), potassium concentrations from 2.3mM to 300mM were added to the wells in triplicates. 6 μl of valinomycin solution from a 250 μM stock solution was added to each well. Then, 3 μl of 3mM DiOC2(3) was added to each well for a final 30 μM concentration in total volume. The volume of each well was made up to 300 μl by adding 50 mM pH 7 HEPES buffer. The plate reader is a filter-based system, and the dye was excited with a 480/30 nm bandpass filter and the emission was using a 530/30 nm and 620/30 nm bandpass filter to measure green and red fluorescence respectively. The R/G was calculated by the device itself. After adding the dye, the fluorescence intensity was measured every 10 minutes and the final reading was recorded after 30 minutes as the signal was stabilized. Data from 6 sets of measurements were obtained. To measure the membrane potential of the cells in the presence of 0 mM, 70 mM, 100 mM, 200 mM, 300 mM, and 400 mM NaCl, the same protocol was followed except for the valinomycin addition.

### Tethered Bead Assay

#### Sample preparation

Tunnel slides were made by layering a square coverslip (22 mm x 22 mm) and rectangular coverslip (60 mm x 24 mm and) between two strips of double sticky tape (fig 1) forming a channel of 24 x 8mm dimension and ∼ 20 μl volume. To immobilize the cells on the glass surface, poly-L-Lysine (0.1% w/v in H_2_O) was added to the flow chamber and incubated for 15 minutes. Unbound poly-L-Lysine were removed by washing with excess of motility buffer (50 μl). The cell suspension (in pH 7 motility buffer) was added to the flow chamber and incubated for 15 minutes. The unbound cells were removed by washing it with pH 7 motility buffer (∼50 μl). Stock solution of polystyrene beads of size 1.1 μm were diluted (1:50 dilution factor) in motility buffer supplemented with 0.002% Tween20. Tween prevented clumping of beads and improved the binding of beads onto the filaments. The bead solution was sonicated for 5 minutes and immediately added to the slide. Adding tween to MB and sonicating the bead solution prevented clumping of beads. The beads were incubated with the cells for 20 minutes. To prevent evaporation, a pool of buffer was kept on both the sides of the slide. After incubation, the unbound beads were washed with ∼ 50 μl of pH 7 motility buffer. Solution exchange was performed by adding solution (of various osmolarities) on one side of the tunnel slide and wicked through the other side with a tissue paper. About ∼10-20 s was taken for complete exchange of the solution and movie was recorded immediately. The slide was stuck on the microscopy stage with a scotch tape to prevent its movement the while solution was exchanged.

**Figure 1.**
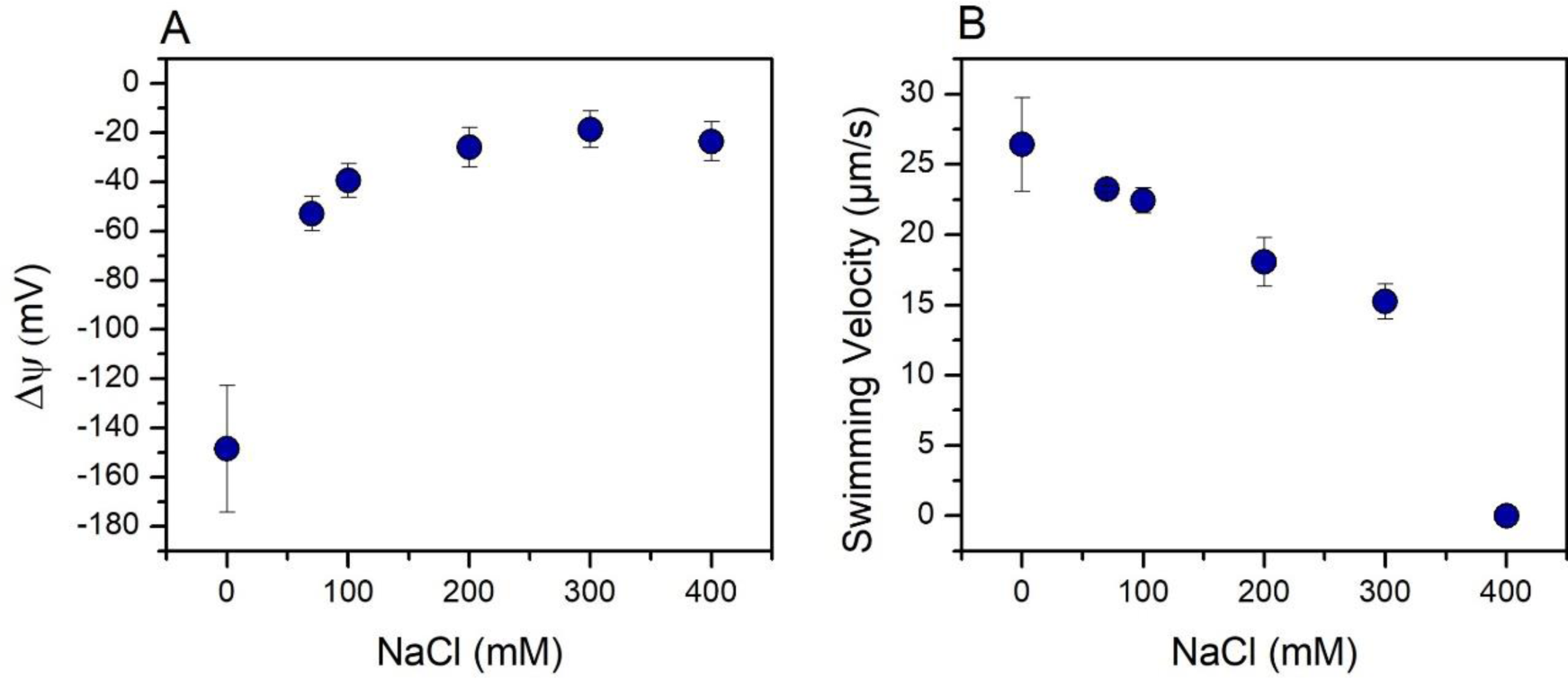
A) Change in membrane potential of *E*.*coli* KAF 84 with NaCl concentration in HEPES buffer. DiOC_2_(3) dye was used to measure the membrane potential in HEPES buffer in a 96 well plate reader. A standard curve of Red/Green fluorescence with change in membrane potential (induced by valinomycin treatment) was made. The curve was not linear at higher and lower membrane potential (see supplementary information) which explains the saturation at higher salt concentration. Nevertheless, the NaCl dissipated the membrane potential. Each data point is mean ± SD. B) Swimming velocity of *Salmonella enteritidis* in pH 7 motility buffer. The cells swam faster in the buffer without NaCl than in the buffer with NaCl. The cells were immotile in 400 mM NaCl. Each data point represents mean of three sets of measurement (∼ 90 cells) ± SD. The experiments were carried out at 30°C.

#### Microscopy and Data acquisition

Tethered bead experiments were carried out with Olympus IX 71 inverted microscope. The microscope is equipped with a 1.4 NA 60X oil immersion objective, sCMOS camera (Andor Xyla), and halogen lamp for bright field illumination. The experiments were carried out at 30°C by fixing the slide on a custom-built temperature controller. Brightfield images of 512×512 pixels was recorded for 20 seconds with 2 ms exposure time. The frame rate was 227.

#### Data Analysis

The images in tiff format were loaded in the ImageJ software. The movies were binarized and the region of rotating beads were cropped. ImageJ’s default thresholding algorithm was used, and the threshold values were adjusted to avoid pixels other than the bead. The cropped movies were saved as tiff stacks. A Matlab program was developed to track the beads and to calculate the frequency of rotation of the beads. The program consisted of the following parts: *a) Particle tracking algorithm:* The algorithm detected the beads using the centroid function of Matlab (Regionprops image processing tool) and the tracked the beads with time. The centroid location of each frame and the corresponding time was generated which was used to calculate the frequency of rotation using a frequency function. *b) Frequency calculation:* The x-y centroid positions of the beads were averaged using a moving mean filter in Matlab with a sliding window size of 227. Each x and y centroid positions were subtracted from the averaged value which created a four-quadrant plot. From the four-quadrant data, the angle (Θ) and change in angle (dΘ) was calculated. Frequency was calculated by using the arctan function, and the phase jumps were fixed by Unwrap function in Matlab.

#### Torque calculation

Torque of the rotating beads was calculated as described before(46). The frictional drag coefficient of the bead attached to the filament (*f*_*b*_) and the filament (*f*_*f*_) was calculated by the following equation:

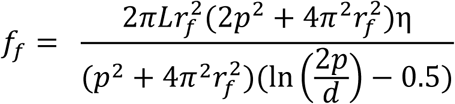

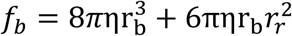

Viscosity of the buffer (η) = 9.6 x 10-4 Ns/m^2^, filament length(L) = 4μm, helical radius (r_f_) = 200nm, helical pitch (p) = 2.3 μm, radius of the filament (d) = 10nm, r_b_ and r_r_ are the radius of the bead and radius of bead rotation respectively. The radius of the bead rotation was obtained from the short axis of the ellipse fit of the centroid position data (See supplemental information Fig.S1). As the motor operates in the low Reynolds number regime, the frictional drag coefficient is the sum of the *f*_*f*_ and *f*_*b*_, and the torque is calculated as:

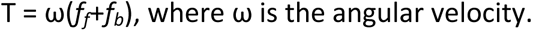

## Results

### a) Effect of NaCl on the electric potential and swimming speed of bacteria in pH 7

We dissipated the electric potential of the cells pH 7 using NaCl and studied its effect on the output of the flagellar motor by analysing the swimming speed of *Salmonella enteritidis*, and the rotational speed and torque of single motors of *E*.*coli* KAF 84. Then, we correlated the switching of flagellar motors to the speed and torque of the motor.

A fluorescent dye based method (see materials and methods) was used to quantify the electric potential of the *E*.*coli* cells in varying concentration of NaCl. Figure 1 A shows that the ΔΨ of the cells decreased with increase in concentration of NaCl in pH 7. The average membrane potential of cells in the absence of NaCl in the buffer was −148.36 ± 25.74 mV and it was decreased to −52.91 ± 6.97 mV in 70 mM NaCl. Average membrane potential values in 100, 200, 300, and 400 mM NaCl was −26.0±7.97 mV, −18.64±7.49 mV, −23.54±7.97 mV respectively. Thus, higher salt concentration dissipated more than 80 % of membrane potential. The average membrane voltage at higher concentration of NaCl (>200mM) was - 22.72 ± 3.73 mV.

The swimming velocity of the cells decreased with increase in NaCl concentration in the motility buffer at pH 7 (Fig.1B). The cells were suspended in the pH 7 motility buffer without NaCl. Before observing the cells at a given NaCl concentration, the cells were washed and resuspended in motility buffer with NaCl. The average velocity of the cells decreased to ∼40 % in 300 mM NaCl. In 400mM NaCl, the cells were immotile. Compared to the swimming velocity in 70 mM NaCl motility buffer, the cells swam ∼ 10% faster in motility buffer without NaCl. The cells regained motility after 30 minutes in 400 mM NaCl although they were tumbly, and when the cell suspension in 400 mM NaCl were washed and resuspended in 70 mM NaCl buffer, they showed vigorous motility (data not shown).

### b) Effect of step-wise increase in NaCl concentration in motility buffer on single motor speed and torque

We observed reduction in motor speed with temporal shifts in NaCl concentration in the motility buffer. The motor speed was observed via a 1.1 μm bead attached to a single flagellar filament (Fig.2A). Unlike the swimming assay, we did not use buffer without NaCl because the bead attachment to the filament requires ionic strength, and we did not obtain tethered beads in the absence of NaCl. Hence, the tethering experiments were carried out in pH 7 motility buffer with 70 mM NaCl. Before recording the first movie (M1), MB solution was added to the tunnel slide. The motors’ mean speed and switching rate decreased with increase in concentration of NaCl in the buffer (Fig.2B). The motors stopped when the concentration of NaCl was shifted to 400mM.

**Figure 2.**
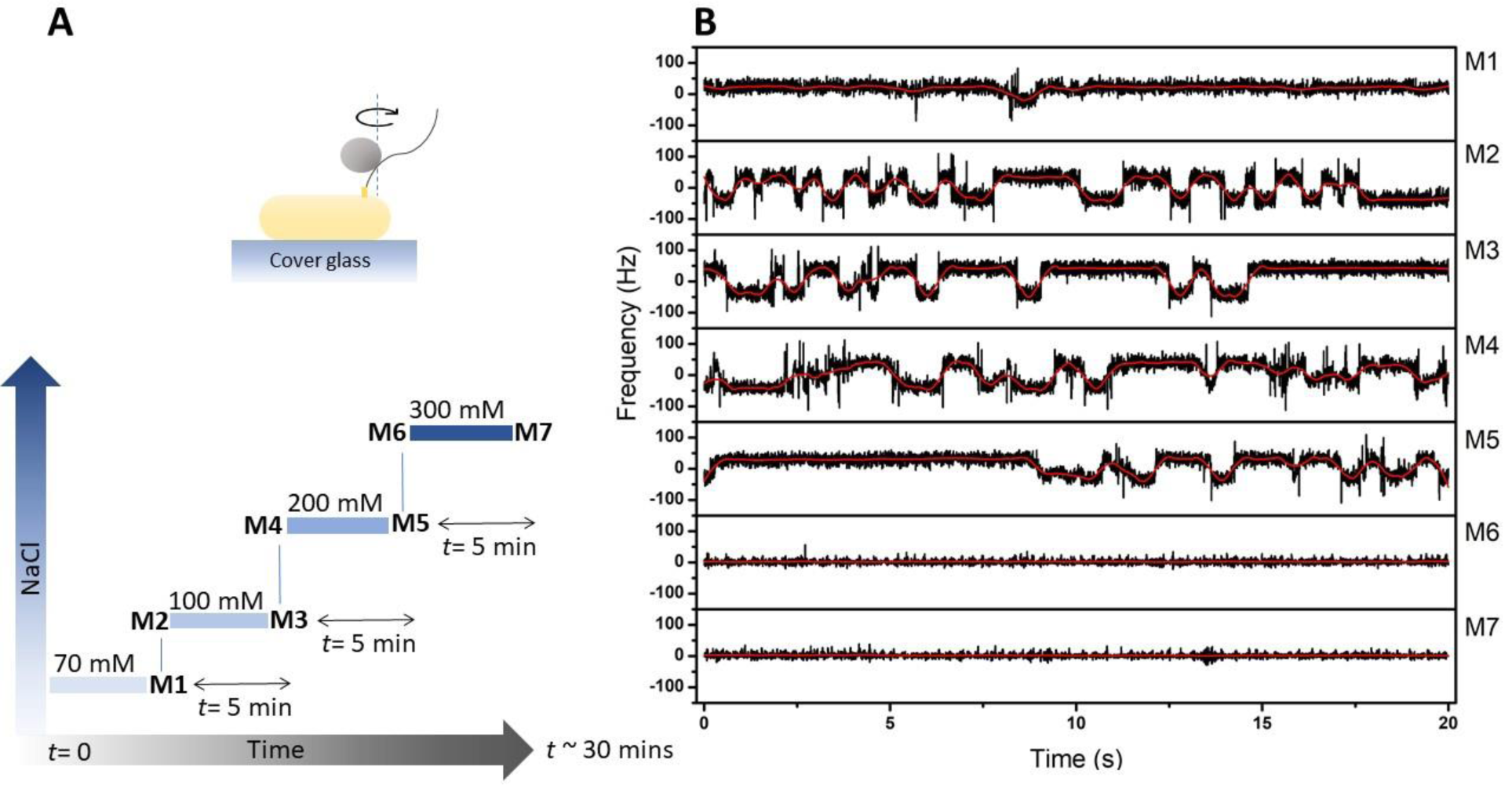
(A) Schematic of the experiment. For each motor, the concentration of NaCl in the motility buffer was increased stepwise by adding motility buffer with different NaCl concentrations. M1-M7 denotes the movies recorded after solution exchange. M1 is the first movie of beads rotating in 70mM NaCl pH 7 motility buffer. M2, M4, and M6 denotes the movie recorded within 30 seconds of exchanging the motility buffer with 100 mM, 200 mM, 300 mM NaCl respectively. M3, M5, M7 denotes the movies recorded after 5 minutes in 100 mM, 200mM, and 300 mM NaCl respectively. B) The frequency of CCW (positive values) and CW rotation (negative values) of 1.1 μm bead attached a flagellar filament in pH 7 motility buffer with various NaCl concentrations and time. 70mM NaCl (M1), 100mM NaCl (M2), 5 minutes in 100mM NaCl (M3), 200mM NaCl (M4), 5 minutes in 200mM NaCl (M5), 300mM NaCl (M6), 5 minutes in 300mM NaCl pH7 motility buffer(M7). The black line represents the raw frequency data and the red line represents the averaged raw frequency. The switching rate of the motor was also decreased with stepwise increase in NaCl concentration in the motility buffer, and the motor was biased to CCW rotation in 300mM NaCl. The data shown is of a single motor.

The motors rotated in both the directions in equal magnitude in all speed range (Fig.3A). However, the magnitude of the torque in both the directions differed (R^2^ = 0.43), perhaps due to the difference in radius of rotation of the beads (Fig.3B). We assumed the frictional drag coefficient of the filament as negligible and only the drag coefficient of the bead was considered. This further explains the variations in the torque with respect to the speed.

**Figure 3.**
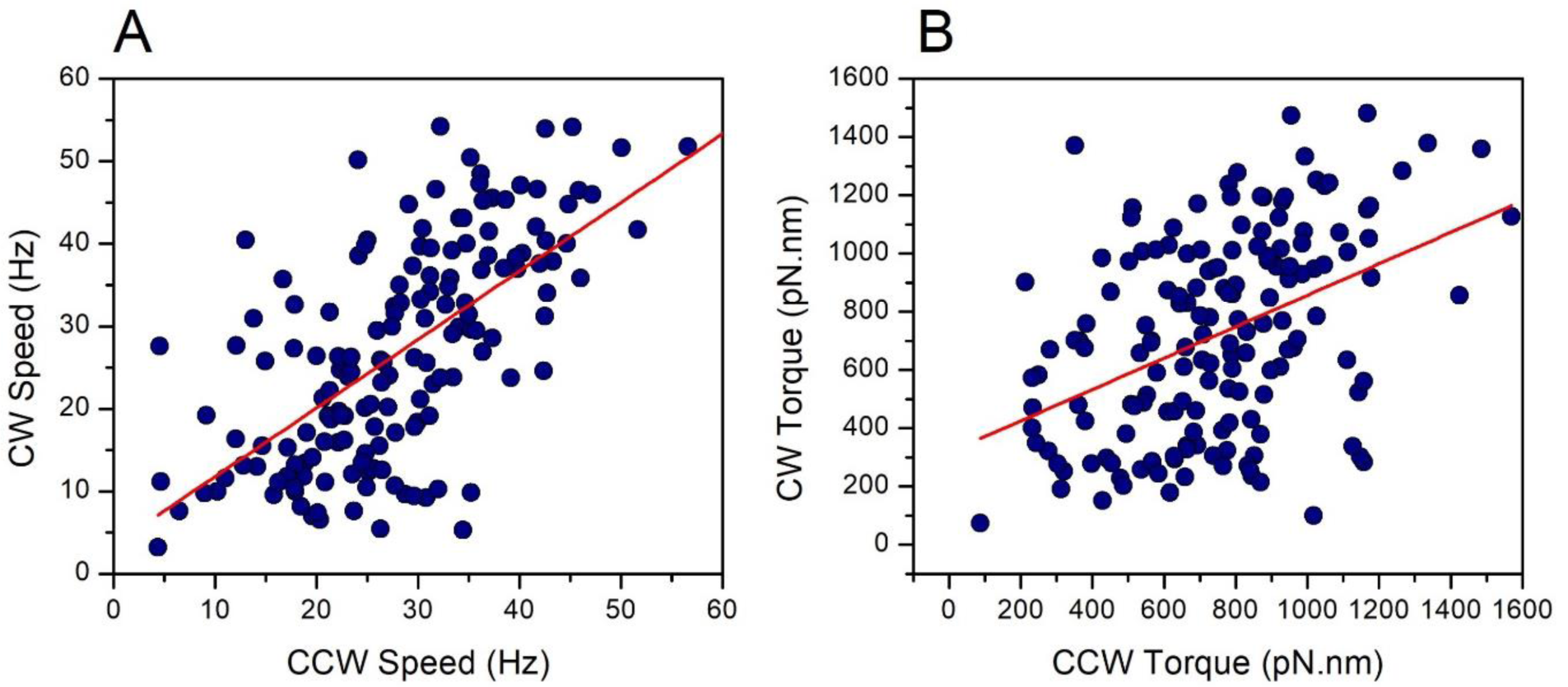
(A) Speed of the flagellar motors with 1.1 μm tethered bead on the filament in pH 7 motility buffer with various NaCl concentrations with linear fit (R^2^ = 0.65). (B) Torque of the motors analysed. Red line is the linear fit (R^2^ = 0.43). The number of data points is 159.

Figure 4 shows the normalised speed of motors analysed. The mean speed of motors in each condition was normalised to the mean of all the motors in 70 mM NaCl. The absolute mean speed of the motors in 70 mM NaCl was 28.57 ± 12.71 Hz and the mean speed of the motors increased 9% when 100 mM NaCl MB was exchanged, and the speed was further increased in the same magnitude after 5 minutes in 100 mM NaCl. The motors did not recover its speed (to the mean speed at 70 mM NaCl) after 5 minutes, neither in CW nor in CCW direction at higher concentration of NaCl (Fig.4). As expected, the trend was similar in both the directions. The speed of the motors reduced to 33 % in 300 mM and rotated at a mean speed of 19.08 ± 1.9 Hz. The motors stopped in 400 mM NaCl and did not recover the speed even after 5 minutes. Also, a downshift to 70mM NaCl (from 300 mM NaCl) stopped most of the motors. The CCW and CW speed at lower speed range varied by more than 50 %, as the motors rotated at a mean CCW speed of 19.08 ± 7.4 Hz in 300 mM NaCl and a mean CW speed of 9.72 ± 2.60 Hz. However, at higher speed range, motors rotated at similar magnitude in both the direction.

**Figure 4.**
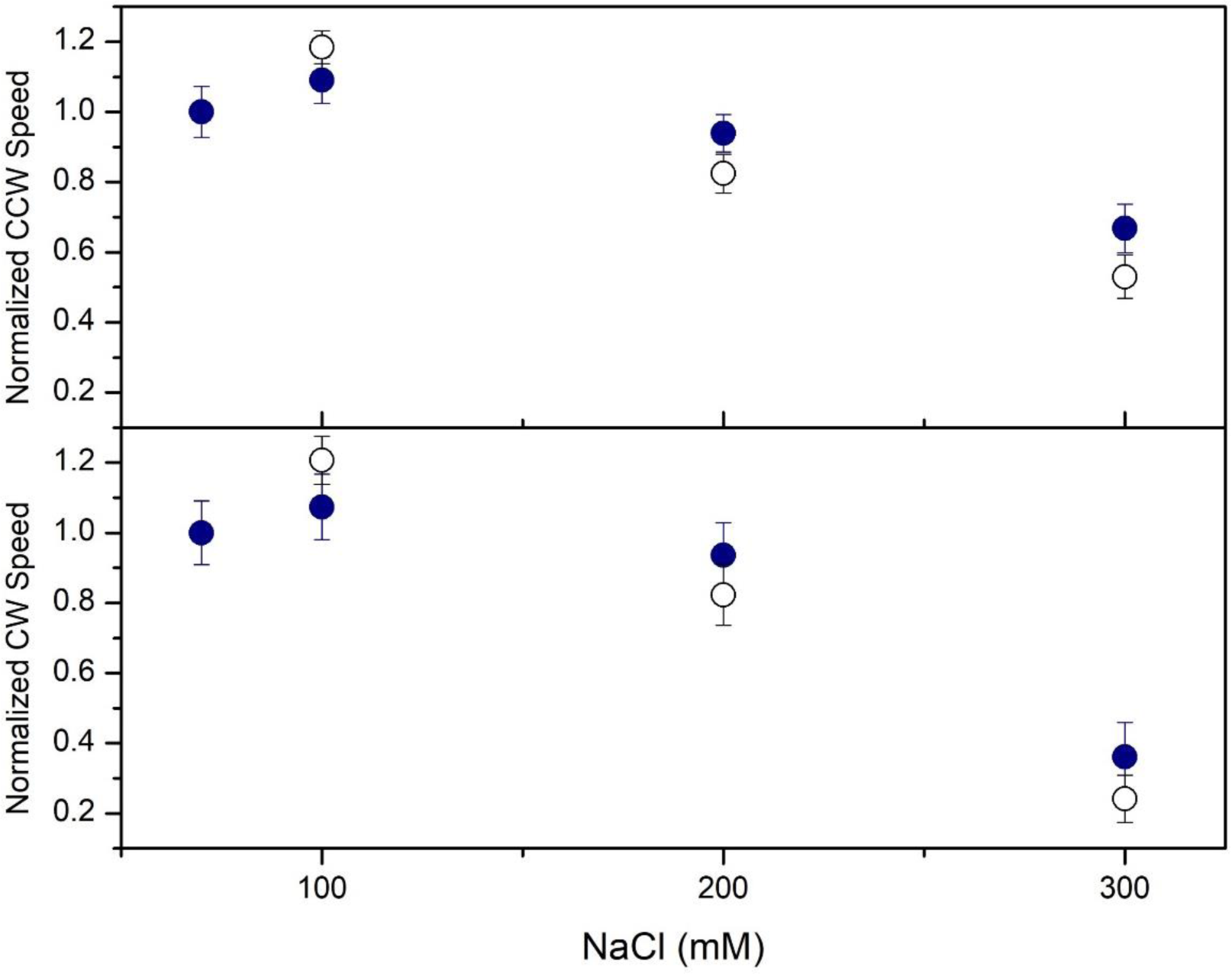
Normalised CCW and CW speed of *E*.*coli*’s flagellar motor with stepwise increase in NaCl concentration. The mean speed in each condition was normalised with the mean speed in 70 mM NaCl to avoid cell to cell variations. Navy blue solid circles represent mean speed after exchanging the solution and open circles represent mean speed after 5 minutes in each NaCl concentration. For each motor, buffer solution was exchanged to increase the concentration of the NaCl stepwise. The motors rotated faster in 100 mM NaCl and decreased in higher concentration of NaCl. The number of cells studied in each condition is 36, 34, 25, 9 for 70mM, 100mM, 200mM, and 300mM NaCl respectively. And the number of cells studied in each condition after 5 minutes were 33, 25, and 12 for 100mM, 200mM and 300mM respectively. Each data point represents mean ± SEM.

As shown the figure 5, the torque changed linearly with speed in both the directions, which was expected as the torque was calculated as a factor of the speed of rotation. The maximum mean torque was 860.81 ± 257 pN.nm in CCW direction. The value in CW direction was similar as the speed was symmetrical. The minimum mean torque was in the CW direction because the slowest rotation of the motors recorded in the CW direction (Fig.5B).

**Figure 5.**
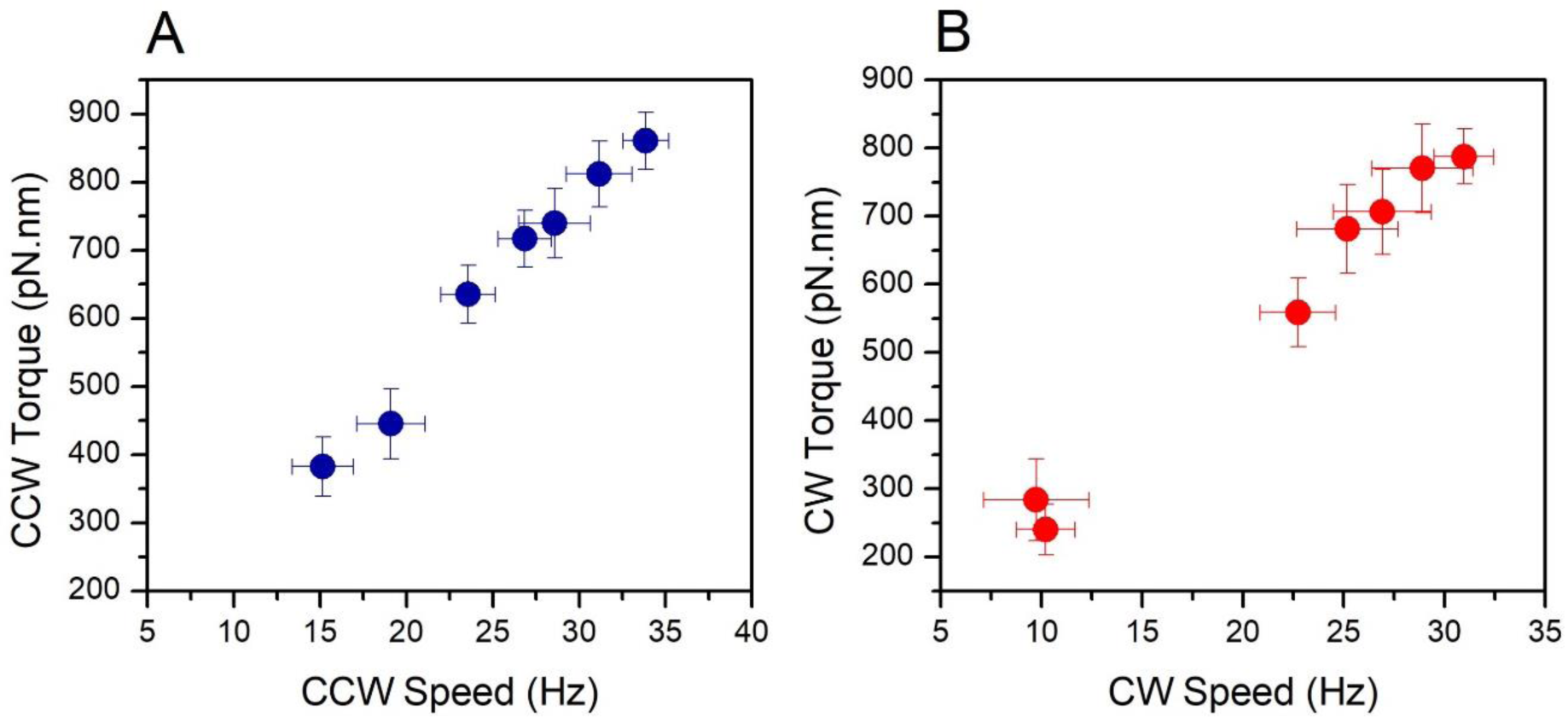
The change in torque with rotation speed of the flagellar motors of *E*.*coli*. The torque was estimated by computing the drag coefficient of spherical bead and frequency of rotation (see materials and methods). The torque changed linearly with speed in both the directions. The radius of rotation of the beads differed among the motor analysed. The lowest speed was accounted in the CW direction (in 300 mM NaCl).

### c) Effect of stepwise increase in NaCl concentration on rotational bias, CW and CCW intervals of motors

Figure 6A and 6B shows the rotational bias of the motor which was calculated as the fraction of time the motors rotated in either direction

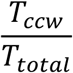

Where T_CCW_ is the time the motors spent in CCW direction and T_total_ is the total duration of the movie (20 seconds). The CW bias was calculated by subtracting the CCW bias from the total time.

**Figure 6.**
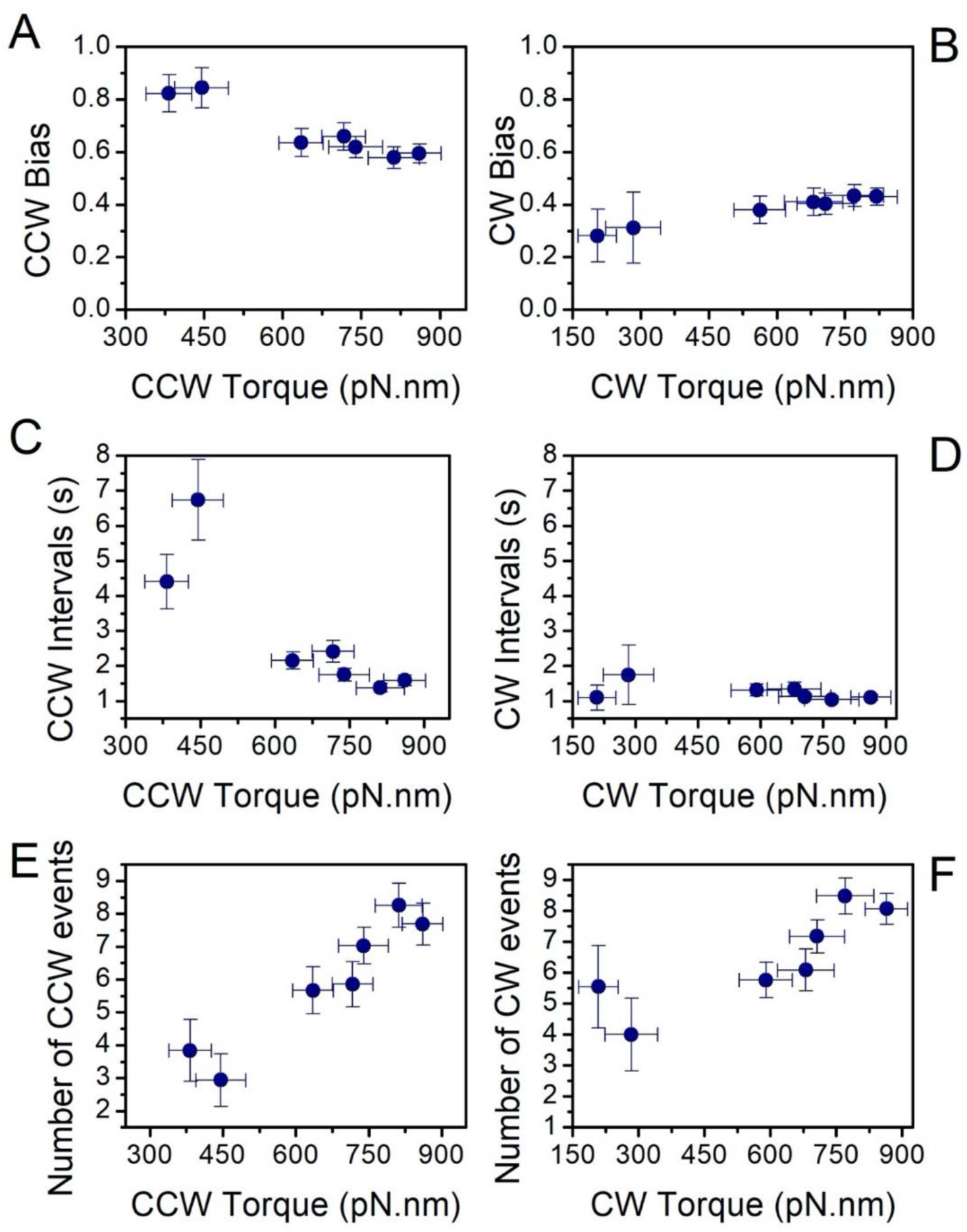
Rotational directionality of motors. (A-B) The change in CCW and CW bias of 1.1 μm bead tethered on flagellar filament in pH 7 motility buffer. The duration of a movie was 20 seconds. The motors were biased to CCW direction at low torque (low speed). At higher torque (higher speed), the motors switched frequently. CW biased motors were not observed in the assay. C-D) Average duration of intervals. The velocity time series was converted to binary series and the intervals were computed for each movie. Change in average duration of intervals with torque in both the directions. The CW duration was constant irrespective of the torque and the average duration in CCW direction was longer in low torque motors. E-F) The change in number of CCW and CW events with torque. As the motor switched more frequently at higher torque, the number of CCW and CW events were similar. As the torque decreased, the number of CCW events decreased while the number of CW events did not change significantly. Each data point represents mean ± SEM.

The motor’s switching was correlated to the changes in torque. In 70 mM NaCl, the CCW bias of the motors were 0.62 ± 0.24, and in 100 mM NaCl, the CCW bias decreased to 0.57 ± 0.24. The CCW bias significantly increased (0.84 ± 0.31) when the concentration increased from 200 mM to 300 mM. Thus, the CCW to CW switching rate decreased with the stepwise increase in NaCl concentration and majority of motors rotated exclusively in CCW rotation in 300mM NaCl (Fig.6A and 6B).

As expected, the average CCW intervals increased with upshock magnitude (Fig.6C and 6D). In 70 mM NaCl, the average CCW interval was 1.74 ± 2.87 s. As the torque of motors increased (due to increase in speed), the mean CCW interval decreased to 1.34 ±2.14 s. As the torque of motors increased, the motors switched frequently (shorter CCW intervals). The slow rotating motors (in 300 mM NaCl) spent an average of 6.7 ± 7.7 seconds in CCW direction (Fig.6C). Correspondingly, the number of CCW events was more as the motors switched more frequently at higher torque, and in 300 mM NaCl concentration, the number of CCW events was less (Fig.6D and 6E).

## Discussion

We used NaCl to dissipate the membrane potential of bacteria and analysed the effect on NaCl concentration in the motility buffer on the speed and rotational bias of *E*.*coli* flagellar motors as well as the speed and tumbling of swimming *Salmonella enteritidis* cells. We observed a reduction in rotational speed of flagellar motor and swimming speed of cells with stepwise increase in NaCl concentration. The motors did not recover the speed 5 minutes post-shock. At higher concentration of NaCl, the motors lost the motility with no recovery. The motors exhibited a CCW rotational bias with increase in concentration of NaCl and the swimming cells were more ‘tumbly’. To study behaviour of motors with abrupt changes in NaCl concentration in the medium, we did not recover the volume of the cell. Thus, the data in this work show the behaviour of individual BFMs and population of swimming cells to abrupt changes in NaCl in the motility buffer.

Membrane potential of bacteria decreases in the presence of NaCl(47). As the proton motive force powers the flagellar motor, the reduction in speed with an increase in concentration of NaCl is due to the dissipation of membrane potential. In pH 7, electric potential is the most dominating component of proton motive, and as pH 7 is the optimal pH (to avoid any non-specific factors affecting the flagellar output) of the bacterial cells used in this study, we performed the experiments in pH 7 motility buffer. In figure 1, the change in speed was not linear with the changes in the membrane potential, contrary to what has been observed before on single motors(14). The membrane potential was found to be saturated above 200 mM NaCl. This is due to the loss of linearity of the dye’s signal with that of the membrane potential, which was evident from the standard curve of the dye (Fig.S4). Fluorescent dyes loses its linearity at higher and lower membrane potential for gram negative bacteria (48). Also, in gram negative bacteria, the response of DiOC_2_(3) dye is not proportional to the proton potential across the cell membrane. Nevertheless, extrapolating the linear trend shows that the membrane potential decreases with increase in NaCl concentration. As the effect of NaCl on the membrane potential has been established before, we aimed to look at the effect of NaCl on membrane potential semi-qualitatively. A plausible explanation for this dissipation of membrane potential is that the cells energetically transports (using the dominant component of the proton motive force) Na^+^ ions into the cell(43). The immotile swimming cells in 400 mM NaCl motility buffer regained motility (with vigorous tumbling, data not shown) after ∼1 hour further supports this argument. Hence, an electrogenic transport of Na^+^ ions into the cytoplasm from the external media causes dissipation of the membrane potential at pH 7.

We observed a stark reduction in swimming speed of *Salmonella enteritidis* and the speed *E*.*coli*’s motors with an increase in concentration of the NaCl in the motility buffer. In MB^-^, the cells swam faster than in the 70 mM NaCl which is usually considered as a control, and the cells were immotile in 400 mM NaCl. We tethered 1.1 μm beads to the flagellum which limits the speed of the motors close to 50 Hz. As we incubated the cells in 70 mM pH 7 motility buffer to tether the beads onto the flagellum, the speed and switching frequency of the motor might have reached a steady state value, and these motors rotated almost exclusively in the CCW (Fig.2A). An interesting thing to note here is that, before recording the first movie (M1), we flowed a fresh 70mM NaCl motility buffer, and the motor’s speed and switching frequency did not change. However, the switching frequency and the speed of the motors increased after we exchanged the medium with 100 mM NaCl motility buffer (Fig.2B). Although the speed remained constant, the switching frequency (CCW-> CW) decreased after 5 minutes in 100 mM NaCl. Further, the addition of 200mM NaCl buffer lowered the motor’s speed and increased the switching frequency (Fig.2D). In 300mM NaCl motility buffer, the speed of the motors reduced significantly, and the motors rotated exclusively in the CCW direction, even after 5 minutes post stimulus (Fig.2F and 2G). To eliminate the variations in motor speed due to changes in metabolic state of cells, we normalised the speed of single motors (Fig.4). Since the load is constant, and the speed of the motors changed only due to the loss of membrane potential, the changes in switching frequency can be attributed to the changes in energy as well. It also suggests that the motor’s switch is sensitive to the changes in speed of the motor.

The reduction in swimming speed at population level and single motors can be explained in three different ways: *1) Dissipation of membrane potential:* NaCl dissipates the membrane potential of the cells, and since the membrane potential powers the motor in pH 7, the motors’ diminished speed is due to the reduction in membrane potential. Since the speed is reduced in both swimming cells and single motors, we emphasize that the cause would be more fundamental like the proton motive force. 2) *High external salt concentration impairs the electrostatic interaction between the rotor and stator:* The flagellar motor is thought to rotate due to the electrostatic interaction between rotor and stator. It has been shown that an increase in ionic strength would impair the motor function as it would interfere with charge residues in the motor(49). The high ionic strength in the external media changes the intracellular ionic strength(50). We have also observed the same effect when NaCl is replaced by KCl in the motility buffer (unpublished data). This supports the previous studies on the motility of wild type and charged mutants in the presence of KCl. Another explanation in this regard is the salt induced misfolding of the motor proteins(49). 3) *Disengagement of stators around the rotor:* At the outset, the stators were thought to be static. However, they were found out to be dynamic as they engage and disengage with the motor due to changes in load, torque, and even ion motive force(3, 4, 20). Recently, studies on *Shewanella oneidensis* MR-1(a bacteria that uses both Na^+^ and H^+^ powered motor) flagellar motor showed that the stators were less incorporated to the motor in the presence of higher Na^+^ concentration in the external media(51, 52). At very high load, such as in tethered cells and motors tethered with 1.1 μm beads, all stators are engaged with the motor which would results in a steady state motor speed. However, the loss of membrane potential uncouples the MotAB stators from the motors(20, 23). This effect is fundamental to the flagellar motor’s mechanism since the ion motive force and the presence of Na^+^ ions affect the assembly of stators around the *Vibrio* motors(44). Thus, the NaCl dissipates the membrane potential which affects the stator incorporation to the motor thereby perturbing the steady state speed of the motors.

The fundamental process of CCW and CW torque generation is symmetrical(53), which explains why the motor instantaneously switches the direction and rotates in the same speed (Fig.3A). Although we estimated the torque from the speed of the motor and drag coefficient of 1.1 μm beads, the changes in torque (Fig.3B) is due to the variations in radius of rotation of the beads. It has been shown that the output of CCW and CW torque varies with load and speed(54), and the rate of switching depends on the motor speed(24, 55). In our study, we observed that the CW motors rotated slower than the CCW motors (Fig.5), which supports these previous studies.

The rate of switching of the motors at a given load changes with the proton motive force and the torque of the motor(56). The switching rate of the motor increased with motor speed (Fig.6A and 6B) and the CCW and CW bias were almost same when motor rotated at maximum speed (70 mM NaCl, high pmf). Previous studies support this observation as well(55). Wang et al observed the motor switching as a non-equilibrium process; that is, the motor’s switch takes input from the motor torque and responds to changes in torque and pmf(56). We observed that the CCW duration was longer than the CW duration at low speed, where the CW duration was constant throughout the range of CW speed (Fig.6C and 6D). The data also means that we did not observe any motor which rotated exclusively in CW direction. Another interesting observation was that the number of CCW and CW events linearly changed with the motor torque in the given direction (Fig.6E and 6F). This means that the number of events in each direction is distributed equally among CCW and CW direction although the speed of the motor varied due to changes in pmf. This shows that the switching of the motor changes with pmf, speed, and even through the osmolytic effect of NaCl.

As sodium chloride is an osmolyte, the behaviour of single motors and the swimming of population of cells can be correlated with osmotaxis. Sodium chloride is a highly permeable osmolyte. Sodium and potassium ions equilibrate across periplasm and cytoplasm within 5 minutes(42). So, we recorded the movies within 30 s of exchanging the buffer and after 5 minutes post-stimulus. Hence, after 5 minutes, the concentration of NaCl will not have any effect on the cell. Osmotically stressed *E*.*coli* exhibits altered metabolism, respiration rate, ATP concentration(57). Rosko et al observed increase in motor speed in the presence of sucrose due to the increase in viscosity of the medium by sucrose(31). Unlike the spatial assay in which the bacteria senses small changes in concentration of osmolytes, in temporal assay the organism faces sudden increase in osmolarity of the medium. We performed the same experiment in pH 8 motility buffer (unpublished data) and observed similar results, suggesting that this process is independent of external pH and it points to the effect of high ionic strength and osmolarity changes of the buffer.

The directionality of the swimming cells was unlike what we saw in single motors. In swimming cell assay, the cells tumbled more with increase in concentration of NaCl (Fig.S3). At 400mM NaCl, the cells lost their motility and recovered the same after 1 hour with more tumble (unpublished data). In 300mM NaCl, the cells tumbled more compared to the lower concentration of NaCl. This may be due to the pseudo-tumbling due osmolarity as described before(30). The individual BFMs showed counterclockwise rotation bias in our experiments. Another explanation for tumbling at higher concentration of NaCl is the higher ionic strength of the buffer affecting the structure of flagellar filaments, which changes the polymorphic transitions of flagella(58, 59). As the shape of the flagellum changes with the presence of salt and pH of the external environment(59), the swimming cells were too ‘tumbly’, which makes the bacterial flagellum inefficient in osmoregulation. Counterclockwise rotation of tethered cells in the presence of high concentration of KCl (0.1-0.6M) was observed by Khan and Macnab(60). In our experiment, the concentration of NaCl is in 0.07 – 0.3M range as we observed loss of motility of individual BFMs and swimming cells in 0.4M NaCl. Also, the effect of the NaCl on the speed of swimming cells was independent of the pH of motility buffer (Fig.S2). The CCW and CW intervals did not follow any distributions. This variability is an attribute of flagellar switching(61–63), and the altered respiration rate and metabolic state of the cells due to changes in osmolarity of the medium.

Studying the effect of NaCl on the flagellar motor is multifaceted: *a)* It helps to look at the mechanism of motor by changing the ion motive force which powers it. *b)* It provides insights into the stator dynamics and the electrostatic mechanism which makes the motor turn. *c)* The mechanism of osmotaxis and how bacteria find optimal water content for its growth and survival. *d)* NaCl’s ability to dissipate the membrane potential makes it suitable to manipulate the pmf to study the mechanism of energy coupling. Although, less is known about the exact mechanism of how NaCl changes the output of the motor, the data in this study paves way into a new direction of flagellar motor research.

## Acknowledgements

This study is funded by Department of Biotechnology, Government of India (Under RGYI scheme to Ravikrishnan E). We thank Nicholas Darnton and Prof Howard Berg of Harvard University for the bead tracking program. The program as used in its original form and no modifications were made. We specially thank Dr. Navish Wadhwa at the Harvard University for providing us the frequency calculation program and for fruitful discussion on developing the bead analysis program used in this study.

## Notes

**Declaration of conflict of interest:** None

## References

1. Costerton JW, Stewart PS, Greenberg EP. 1999. Bacterial Biofilms: A Common Cause of Persistent Infections. Science (80-) 284:1318–1322.

2. Josenhans C, Suerbaum S. 2002. The role of motility as a virulence factor in bacteria. Int J Med Microbiol. Urban & Fischer.

3. Tipping MJ, Delalez NJ, Lim R, Berry RM, Armitage JP, Bassler B. 2013. Load-Dependent Assembly of the Bacterial Flagellar Motor.

4. Lele PP, Hosu BG, Berg HC. 2013. Dynamics of mechanosensing in the bacterial flagellar motor. Proc Natl Acad Sci 110:11839–11844.

5. Larsen SH, Adler J, Gargus JJ, Hogg RW. 1974. Chemomechanical coupling without ATP: the source of energy for motility and chemotaxis in bacteria. Proc Natl Acad Sci U S A 71:1239–43.

6. Matsuura S, Shioi JI, Imae Y, Iida S. 1979. Characterization of the Bacillus subtilis motile system driven by an artificially created proton motive force. J Bacteriol 140:28–36.

7. Shioi JI, Imae Y, Oosawa F. 1978. Protonmotive force and motility of Bacillus subtilis. J Bacteriol 133:1083–8.

8. Shioi JI, Matsuura S, Imae Y. 1980. Quantitative measurements of proton motive force and motility in Bacillus subtilis. J Bacteriol 144:891–7.

9. Miller JB, Koshland DE. 1977. Sensory electrophysiology of bacteria: relationship of the membrane potential to motility and chemotaxis in Bacillus subtilis. Proc Natl Acad Sci U S A 74:4752–6.

10. Glagolev AN, Skulachev VP. 1978. The proton pump is a molecular engine of motile bacteria. Nature 272:280–2.

11. Manson MD, Tedesco P, Berg HC, Harold FM, Van der Drift C. 1977. A protonmotive force drives bacterial flagella. Proc Natl Acad Sci 74:3060–3064.

12. Mitchell P, Moyle J. 1967. Chemiosmotic Hypothesis of Oxidative Phosphorylation. Nature 213:137–139.

13. Shioi J, Imae Y, Matsuura S, After A. 1977. Motility in 82:187–190.

14. Gabel C V, Berg HC. 2003. The speed of the flagellar rotary motor of Escherichia coli varies linearly with protonmotive force. Proc Natl Acad Sci U S A 100:8748–51.

15. Kami-ike N, Kudo S, Hotani H. 1991. Rapid changes in flagellar rotation induced by external electric pulses. Biophys J 60:1350–5.

16. Khan S, Dapice M, Humayun I. 1990. Energy transduction in the bacterial flagellar motor Effects of load and pH Motility measurements 57:779–796.

17. Repaske DR, Adler J. 1981. Change in intracellular pH of Escherichia coli mediates the chemotactic response to certain attractants and repellents. J Bacteriol 145:1196–208.

18. Kihara M, Macnab RM. 1981. Cytoplasmic pH mediated pH taxis and weak-acid repellent taxis of bacteria. J Bacteriol.

19. Minamino T, Imae Y, Oosawa F, Kobayashi Y, Oosawa K. 2003. Effect of Intracellular pH on Rotational Speed of Bacterial Flagellar Motors. J Bacteriol 185:1190–1194.

20. Tipping MJ, Steel BC, Delalez NJ, Berry RM, Armitage JP. 2013. Quantification of flagellar motor stator dynamics through in vivo proton-motive force control. Mol Microbiol 87:338–347.

21. Armitage JP, Evans MCW. 1985. Control of the protonmotive force in Rhodopseudomonas sphaeroides in the light and dark and its effect on the initiation of flagellar rotation. BBA - Bioenerg 806:42–55.

22. Sowa Y, Rowe AD, Leake MC, Yakushi T, Homma M, Ishijima A, Berry RM. 2005. Direct observation of steps in rotation of the bacterial flagellar motor. Nature 437:916–919.

23. Fung DC, Berg HC. 1995. Powering the flagellar motor of Escherichia coli with an external voltage source. Nature 375:809–812.

24. Fahrner KA, Ryu WS, Berg HC. 2003. Bacterial flagellar switching under load. Nature 423:938–938.

25. Krell T, Lacal J, Muñoz-Martínez F, Reyes-Darias JA, Cadirci BH, García-Fontana C, Ramos JL. 2011. Diversity at its best: bacterial taxis. Environ Microbiol 13:1115–1124.

26. Wadhams GH, Armitage JP. 2004. Making sense of it all: bacterial chemotaxis. Nat Rev Mol Cell Biol 5:1024–1037.

27. Blair DF. 1995. How Bacteria Sense and Swim. Annu Rev Microbiol 49:489–522.

28. Bi S, Lai L. 2015. Bacterial chemoreceptors and chemoeffectors. Cell Mol Life Sci. Birkhauser Verlag AG.

29. Massart J. 1889. Sensibilité et adaptation des organismes à la concentration des solutions salinesArch Biol.

30. Adler J, Li C, Boileau AJ, Qi Y, Kung C. 1988. Osmotaxis in Escherichia coli, p. 19–22. In Cold Spring Harbor Symposia on Quantitative Biology. National Academy of Sciences.

31. Rosko J, Martinez V, Poon W, Pilizota T. 2017. Osmotaxis in Escherichia coli through changes in motor speed. Proc Natl Acad Sci U S A 114:E7969–E7976.

32. Li C, Adler J. 1993. Escherichia coli shows two types of behavioral responses to osmotic upshift. J Bacteriol 175:2564–2567.

33. Pilizota T, Shaevitz JW. 2013. Plasmolysis and Cell Shape Depend on Solute Outer-Membrane Permeability during Hyperosmotic Shock in E. coli. Biophys J 104:2733–2742.

34. Korber DR, Choi A, Wolfaardt GM, Caldwell DE. 1996. Bacterial plasmolysis as a physical indicator of viability. Appl Environ Microbiol 62:3939–47.

35. Sleator RD, Hill C. 2002. Bacterial osmoadaptation: the role of osmolytes in bacterial stress and virulence. FEMS Microbiol Rev 26:49–71.

36. Le Rudulier D, Strom AR, Dandekar AM, Smith LT, Valentine RC. 1984. Molecular Biology of Osmoregulation. Science (80-) 224:1064–1068.

37. Miller KJ, Reinhold VN, Weissborn AC, Kennedy EP. 1987. Cyclic glucans produced by Agrobacterium tumefaciens are substituted with sn-1-phosphoglycerol residues. Biochim Biophys Acta - Biomembr 901:112–118.

38. Epstein W. 1986. Osmoregulation by potassium transport in Escherichia coli. FEMS Microbiol Lett 39:73–78.

39. Garrett S, Taylor RK, Silhavy TJ, Berman ML. 1985. Isolation and characterization of delta ompB strains of Escherichia coli by a general method based on gene fusions. J Bacteriol 162:840–4.

40. Vaknin A, Berg HC. 2006. Osmotic stress mechanically perturbs chemoreceptors in Escherichia coli. Proc Natl Acad Sci 103:592–596.

41. Decad GM, Nikaido H. 1976. Outer membrane of gram-negative bacteria. XII. Molecular-sieving function of cell wall. J Bacteriol 128:325–36.

42. Cowie DB, Roberts RB, Roberts IZ. 1949. Potassium metabolism in Escherichia coli. I. Permeability to sodium and potassium ions. J Cell Comp Physiol 34:243–257.

43. Barker SL, Kashket ER. 1977. Effects of sodium ions on the electrical and pH gradients across the membrane of streptococcus lactis cells. J Supramol Struct 6:383–388.

44. Fukuoka H, Wada T, Kojima S, Ishijima A, Homma M. 2009. Sodium-dependent dynamic assembly of membrane complexes in sodium-driven flagellar motors. Mol Microbiol 71:825–835.

45. Pottash AE, McKay R, Virgile CR, Ueda H, Bentley WE. 2017. TumbleScore: Run and tumble analysis for low frame-rate motility videos. Biotechniques 62.

46. Che Y-S, Nakamura S, Kojima S, Kami-ike N, Namba K, Minamino T. 2008. Suppressor analysis of the MotB(D33E) mutation to probe bacterial flagellar motor dynamics coupled with proton translocation. J Bacteriol 190:6660–7.

47. Barker SL, Kashket ER. 1977. Effects of sodium ions on the electrical and pH gradients across the membrane of streptococcus lactis cells. J Supramol Struct 6:383–388.

48. te Winkel JD, Gray DA, Seistrup KH, Hamoen LW, Strahl H. 2016. Analysis of antimicrobial-triggered membrane depolarization using voltage sensitive dyes. ront Cell Dev Biol 4:29.

49. Zhou J, Lloyd SA, Blair DF. 1998. Electrostatic interactions between rotor and stator in the bacterial flagellar motor. Proc Natl Acad Sci U S A 95:6436–41.

50. Kohno T, Roth J. 1979. Electrolyte effects on the activity of mutant enzymes in vivo and in vitro. Biochemistry 18:1386–1392.

51. Paulick A, Koerdt A, Lassak J, Huntley S, Wilms I, Narberhaus F, Thormann KM. 2009. Two different stator systems drive a single polar flagellum in Shewanella oneidensis MR-1. Mol Microbiol 71:836–850.

52. Paulick A, Delalez NJ, Brenzinger S, Steel BC, Berry RM, Armitage JP, Thormann KM. 2015. Dual stator dynamics in the Shewanella oneidensisMR-1 flagellar motor. ol Microbiol 96:993–1001.

53. Nakamura S, Kami-ike N, Yokota JIP, Minamino T, Namba K. 2010. Evidence for symmetry in the elementary process of bidirectional torque generation by the bacterial flagellar motor. Proc Natl Acad Sci U S A 107:17616–17620.

54. Yuan J, Fahrner KA, Turner L, Berg HC. 2010. Asymmetry in the clockwise and counterclockwise rotation of the bacterial flagellar motor. Proc Natl Acad Sci U S A 107:12846–9.

55. Yuan J, Fahrner KA, Berg HC. 2009. Switching of the Bacterial Flagellar Motor Near Zero Load. J Mol Biol 390:394–400.

56. Wang F, Shi H, He R, Wang R, Zhang R, Yuan J. 2017. Non-equilibrium effect in the allosteric regulation of the bacterial flagellar switch. Nat Phys 13:710–714.

57. Wood JM. 1999. Osmosensing by bacteria: signals and membrane-based sensors. Microbiol Mol Biol Rev 63:230–62.

58. Berg HC. 2001. E.Coli in motionSpringer.

59. Kamiya R, Hotani H, Asakura S. 1982. Polymorphic transition in bacterial flagella. Symp Soc Exp Biol 35:53–76.

60. Khan S, Macnab RM. 1980. The steady-state counterclockwise/clockwise ratio of bacterial flagellar motors is regulated by protonmotive force. J Mol Biol 138:563–597.

61. Waite AJ, Frankel NW, Emonet T. 2018. Behavioral Variability and Phenotypic Diversity in Bacterial Chemotaxis. Annu Rev Biophys 47:595–616.

62. Korobkova E, Emonet T, Vilar JMG, Shimizu TS, Cluzel P. 2004. From molecular noise to behavioural variability in a single bacterium. Nature 428:574–578.

63. Park H, Oikonomou P, Guet CC, Cluzel P. 2011. Noise Underlies Switching Behavior of the Bacterial Flagellum. Biophys J 101:2336–2340.

